# DUBStepR: correlation-based feature selection for clustering single-cell RNA sequencing data

**DOI:** 10.1101/2020.10.07.330563

**Authors:** Bobby Ranjan, Wenjie Sun, Jinyu Park, Kunal Mishra, Ronald Xie, Fatemeh Alipour, Vipul Singhal, Florian Schmidt, Ignasius Joanito, Nirmala Arul Rayan, Michelle Gek Liang Lim, Shyam Prabhakar

## Abstract

Feature selection (marker gene selection) is widely believed to improve clustering accuracy, and is thus a key component of single cell clustering pipelines. However, we found that the performance of existing feature selection methods was inconsistent across benchmark datasets, and occasionally even worse than without feature selection. Moreover, existing methods ignored information contained in gene-gene correlations. We therefore developed DUBStepR (<u>D</u>etermining the <u>U</u>nderlying <u>B</u>asis using <u>Step</u>wise <u>R</u>egression), a feature selection algorithm that leverages gene-gene correlations with a novel measure of inhomogeneity in feature space, termed the Density Index (DI). Despite selecting a relatively small number of genes, DUBStepR substantially outperformed existing single-cell feature selection methods across diverse clustering benchmarks. In a published scRNA-seq dataset from sorted monocytes, DUBStepR sensitively detected a rare and previously invisible population of contaminating basophils. DUBStepR is scalable to over a million cells, and can be straightforwardly applied to other data types such as single-cell ATAC-seq. We propose DUBStepR as a general-purpose feature selection solution for accurately clustering single-cell data.

## Introduction

Heterogeneity in single-cell RNA sequencing (scRNA-seq) datasets is frequently characterized by identifying cell clusters in gene expression space, wherein each cluster represents a distinct cell type or cell state. In particular, numerous studies have used unsupervised clustering to discover novel cell populations in heterogeneous samples (1). The steps involved in unsupervised clustering of scRNA-seq data have been well documented (2). i) Low-quality cells are first discarded in a quality control step (3). ii) Reads obtained from the remaining cells are then normalized to remove the influence of technical effects, while preserving true biological variation (4). iii) After normalization, feature selection is performed to select the subset of genes that are informative for clustering, iv) which are then typically reduced to a small number of dimensions using Principal Component Analysis (PCA) (5). v) In the reduced principal component (PC) space, cells are clustered based on their distance from one another (typically, Euclidean distance), and vi) the corresponding clusters are assigned a cell type or state label based on the known functions of their differentially expressed (DE) genes (6). Although feature selection is a critical step in the canonical clustering workflow described above, only a few differ-ent approaches have been developed in this space. Moreover, there have been only a handful of systematic benchmarking studies of scRNA-seq feature selection methods (7–9). A good feature selection algorithm is one that selects cell-type-specific (DE) genes as features, and rejects the remaining genes. More importantly, the algorithm should select features that optimize the separation between biologically distinct cell clusters. A comprehensive benchmarking study of feature selection methods would ideally use both of these metrics.

The most widely used approach for feature selection is mean-variance modeling: genes whose variation across cells exceeds a data-derived null model are selected as features (10, 11). Such genes are described as highly variable genes (HVGs) (12). Some earlier single cell studies instead selected genes with high loading on the top principal components of the gene expression matrix (high loading genes, or HLGs) as features (13). M3Drop, a more recent method, selects genes whose dropout rate (number of cells in which the gene is undetected) exceeds that of other genes with the same mean expression (9). As an alternative approach to detect rare cell types, GiniClust uses a modified Gini index to identify genes whose expression is concentrated in a relatively small number of cells (14). All of the above feature selection methods test genes individually, without considering expression relationships between genes. Another drawback is that existing methods for determining the size of the feature set do not bear direct relation to the separation of cells in the resulting space.

Here, we present <u>D</u>etermining the <u>U</u>nderlying <u>B</u>asis using <u>Step</u>wise <u>R</u>egression (DUBStepR), an algorithm for feature selection based on gene-gene correlations. A key feature of DUBStepR is the use of a step-wise approach to identify an initial core set of genes that most strongly represent coherent expression variation in the dataset. Uniquely, DUBStepR defines a novel graph-based measure of cell aggregation in the feature space, and uses this measure to optimize the number of features. We benchmark DUBStepR against 6 commonly used feature selection algorithms on datasets from 4 different scRNA-seq protocols (10x Genomics, Drop-Seq, CEL-Seq2 and Smart-Seq2) and show that it substantially outperforms other methods. We used DUBStepR to detect a rare, contaminating basophil population in FACS-purified monocytes and dendritic cells (16). Finally, we show that DUBStepR could potentially be applied even to single-cell ATAC sequencing data.

## Results

### Gene-gene correlations predict cell-type-specific DE genes

The first step in DUBStepR is to select an initial set of candidate features based on known properties of cell-type-specific DE genes (marker genes). DE genes specific to the same cell types would tend to be highly correlated with each other, whereas those specific to distinct cell types are likely to be anti-correlated (Figure 1a-b; Methods). In contrast, non-DE genes are likely to be only weakly correlated (Figure 1c). We therefore hypothesized that a correlation range score derived from the difference between the strongest positive and strongest negative correlation coefficients of a gene (Methods), would be substantially elevated among DE genes. Indeed, we found that the correlation range score was significantly higher for DE genes relative to non-DE genes (Figure 1d). Moreover, the correlation range score of a gene was highly predictive of its greatest fold-change between cell types, and also its most significant differential expression q-value (Figure 1e-f). Due to the strong association between correlation range and marker gene status, DUBStepR selects genes with high correlation range score as the initial set of candidate feature genes (Methods).

**Fig.1.**
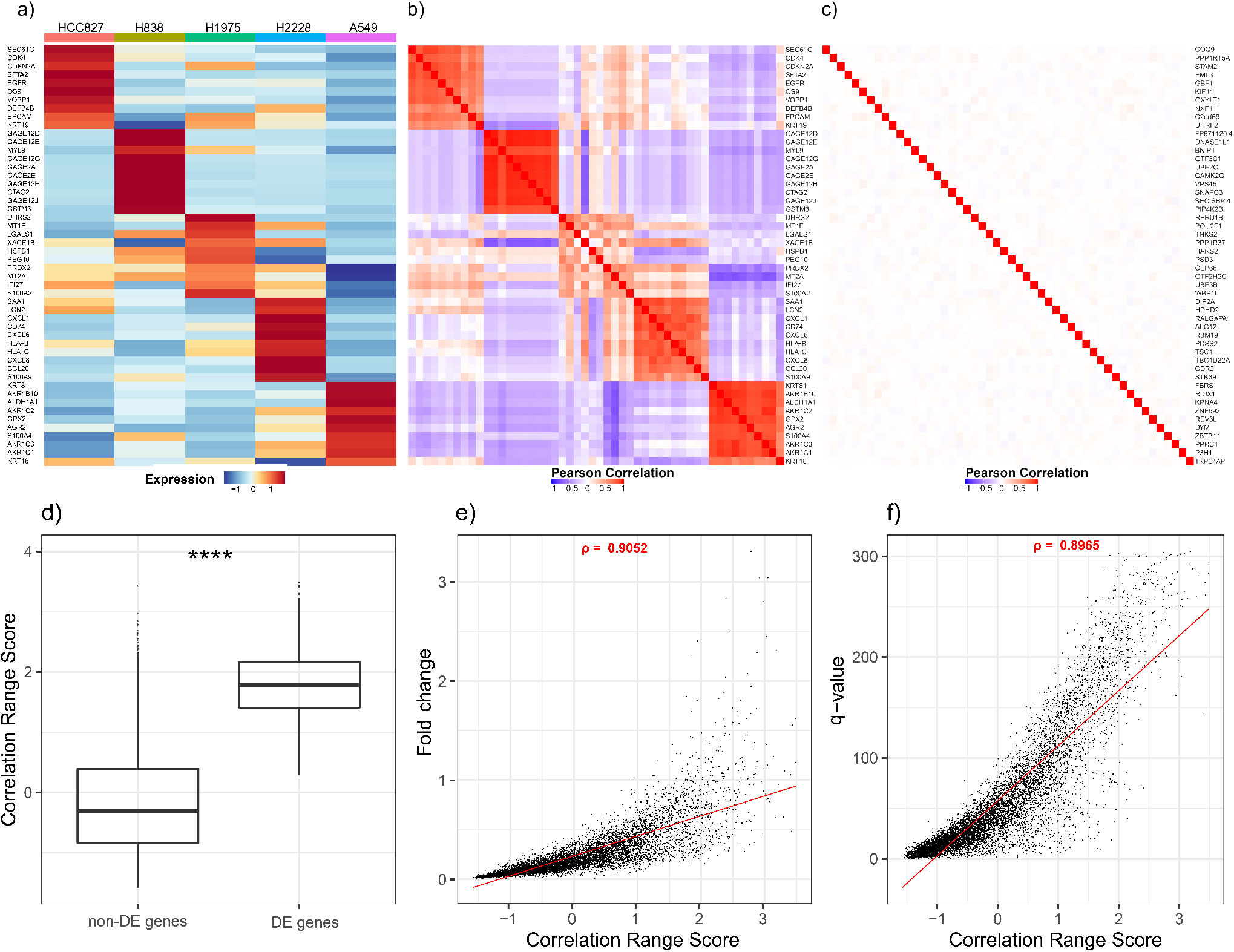
Expression correlations of DE genes: scRNA-seq data from 5 lung adenocarcinoma cell lines (15). a) Average expression of top 10 DE genes for each cell type. b) Gene-gene correlations of the same genes. c) Gene-gene correlations for non-DE genes. d) Boxplot showing correlation range scores for non-DE and DE genes. e-f) Scatter plot of genes showing correlation between e) log2(fold change) of cell-type-specific expression and f) -log10(q-value) of cell-type-specific expression with correlation range score. ****: p-value <= 0.0001. *ρ*: Spearman Correlation.

### Stepwise regression identifies a minimally redundant feature subset

We observed that candidate feature genes formed correlated blocks of varying size in the gene-gene correlation (GGC) matrix (Figure 2a), with each block presumably representing a distinct pattern of expression variation across the cells. To ensure more even representation of the diverse expression signatures within the candidate feature set, we sought to identify a representative minimally-redundant subset, which we termed “seed” genes. For this purpose, DUBStepR performs stepwise regression on the GGC matrix, regressing out, at each step, the gene explaining the largest amount of variance in the residual from the previous step (Figure 2b-d). We devised an efficient implementation of this procedure that requires only a single matrix multiplication at each step (Methods).

**Fig.2.**
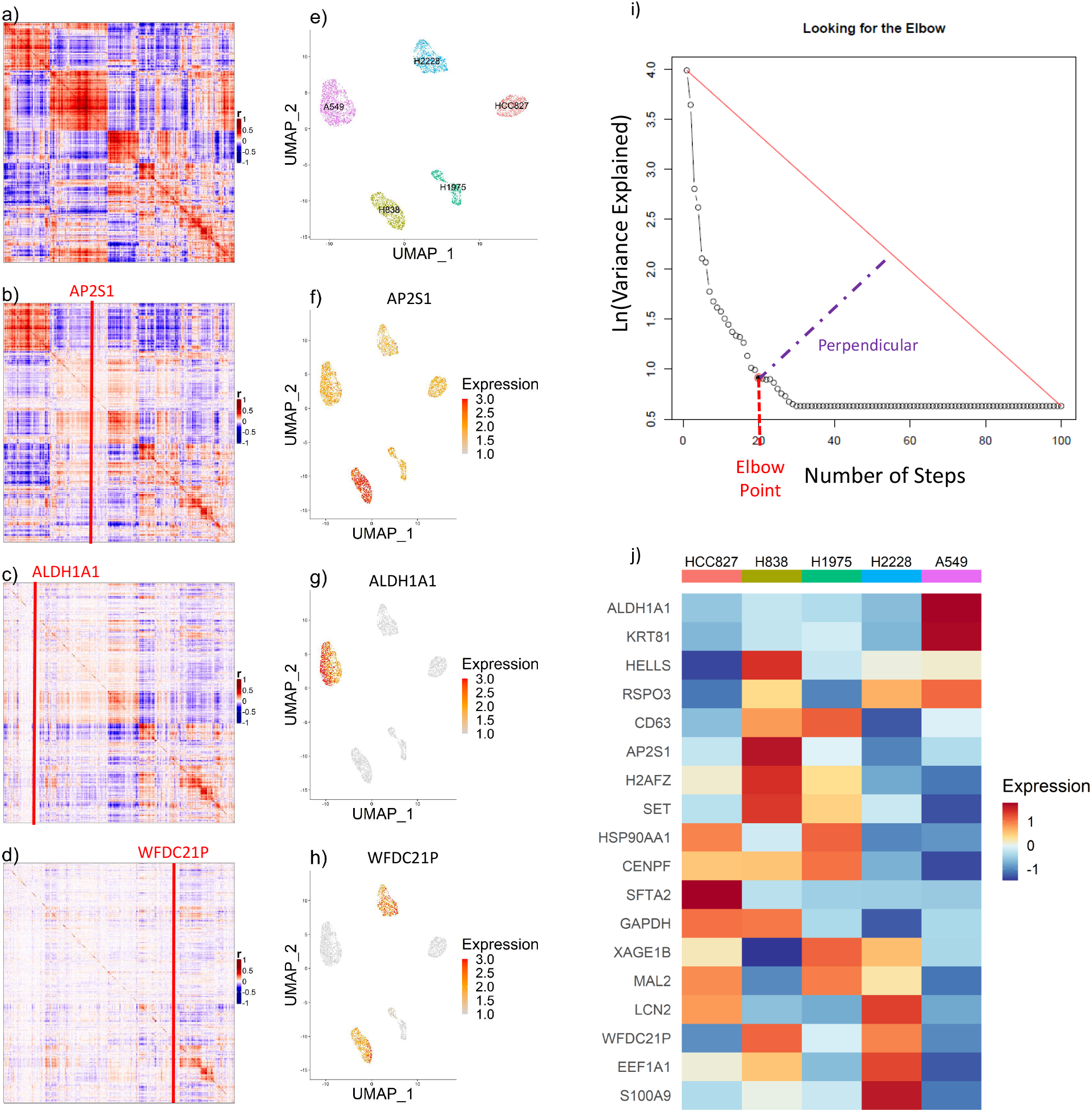
a) Gene-gene correlation matrix of candidate feature genes (high correlation range score). b-d) Residuals from stepwise regression on the gene-gene correlation matrix. e) UMAP visualization of cells in an optimal feature space, colored by cell line. f-h) Same UMAP, colored by expression of genes regressed out in the first 3 steps. i) Scree plot: variance in GGC matrix explained by the gene regressed out at each step. j) Standardized average expression of the final seed gene set in each of the 5 cell lines.

This approach selects seed genes with diverse patterns of cell-type-specificity (Figure 2e-h). DUBStepR then uses the elbow point of the stepwise regression scree plot to determine the optimal number of steps (Methods), i.e. the size of the seed gene set (Figure 2i,j).

### Guilt-by-association expands the feature set

Although the seed genes in principle span the major expression signatures in the dataset, each individual signature is now represented by only a handful of genes (2-5 genes, in most cases). Given the high level of noise in scRNA-seq data, it is likely that this is insufficient to fully capture coherent variation across cells. DUBStepR therefore expands the seed gene set by iteratively adding correlated genes from the candidate feature set (Supp. Fig. S4; Methods). This process is continued until DUBStepR reaches the optimal number of feature genes (see below).

### Benchmarking

To benchmark the performance of DUB-StepR, we compared it against 5 other algorithms for feature selection in scRNA-seq data: three variants of the HVG approach (HVGDisp, HVGVST, trendVar), HLG and M3Drop/DANB (Table 1). For completeness we also bench-marked GiniClust, though it was designed only for identifying markers of rare cell types. Each algorithm was bench-marked on 7 datasets spanning 4 scRNA-seq protocols: 10X Genomics, Drop-Seq, CEL-Seq2, and Smart-Seq on the Fluidigm C1 (Supplementary Note 2A). These datasets were selected because the true cell type could be independently ascertained based on cell line identity or FACS gate. Our benchmarking approach thus avoids the circularity of using algorithmically defined cell type labels as ground truth.

**Table 1.**
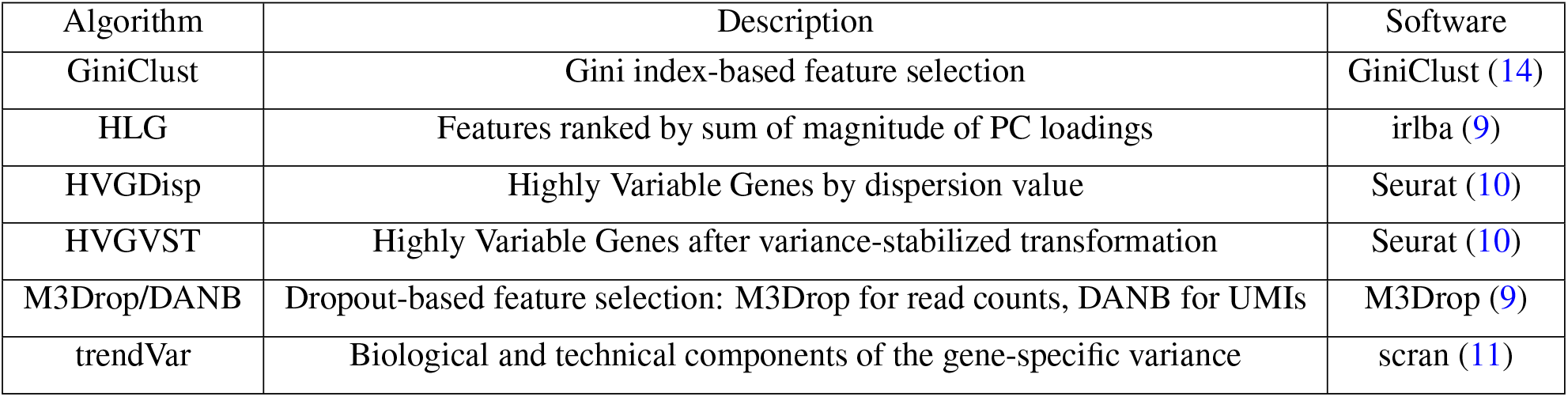
Feature selection methods used for the benchmarking comparison.

To evaluate the quality of the selected features, we used the well-established Silhouette Index (SI), which quantifies cluster separation, i.e. closeness between cells belonging to the same cluster, relative to distance to cells from other clusters (17). In addition to being a well established measure of single cell cluster separation (18) (19) (20), the SI has the advantage of being independent of any downstream clustering algorithm. We evaluated the SI of each algorithm across a range of feature set sizes (50-4, 000), scaled the SI values to a maximum of 1 for each dataset and then averaged the scaled SIs across the 7 datasets (Figure 3a; Supplementary Figure S2). Remarkably, HLG, an elementary PCA-based method that predates scRNA-seq technology, achieved greater average cell type separation than existing single cell algorithms at most feature set sizes. M3Drop, HVGDisp and trendVAR remained close to their respective performance peaks over a broad range from 200 to 2, 000 features, and dropped off on either side of this range. HVGVST, GiniClust and HLG showed a more peaked performance characteristic, with maxima at 500-1, 000 features. DUBStepR substantially outper-formed all other methods across the entire range of feature set size. Moreover, DUBStepR was the top-ranked algorithm on 5 of the 7 datasets (Figure 3b).

**Fig.3.**
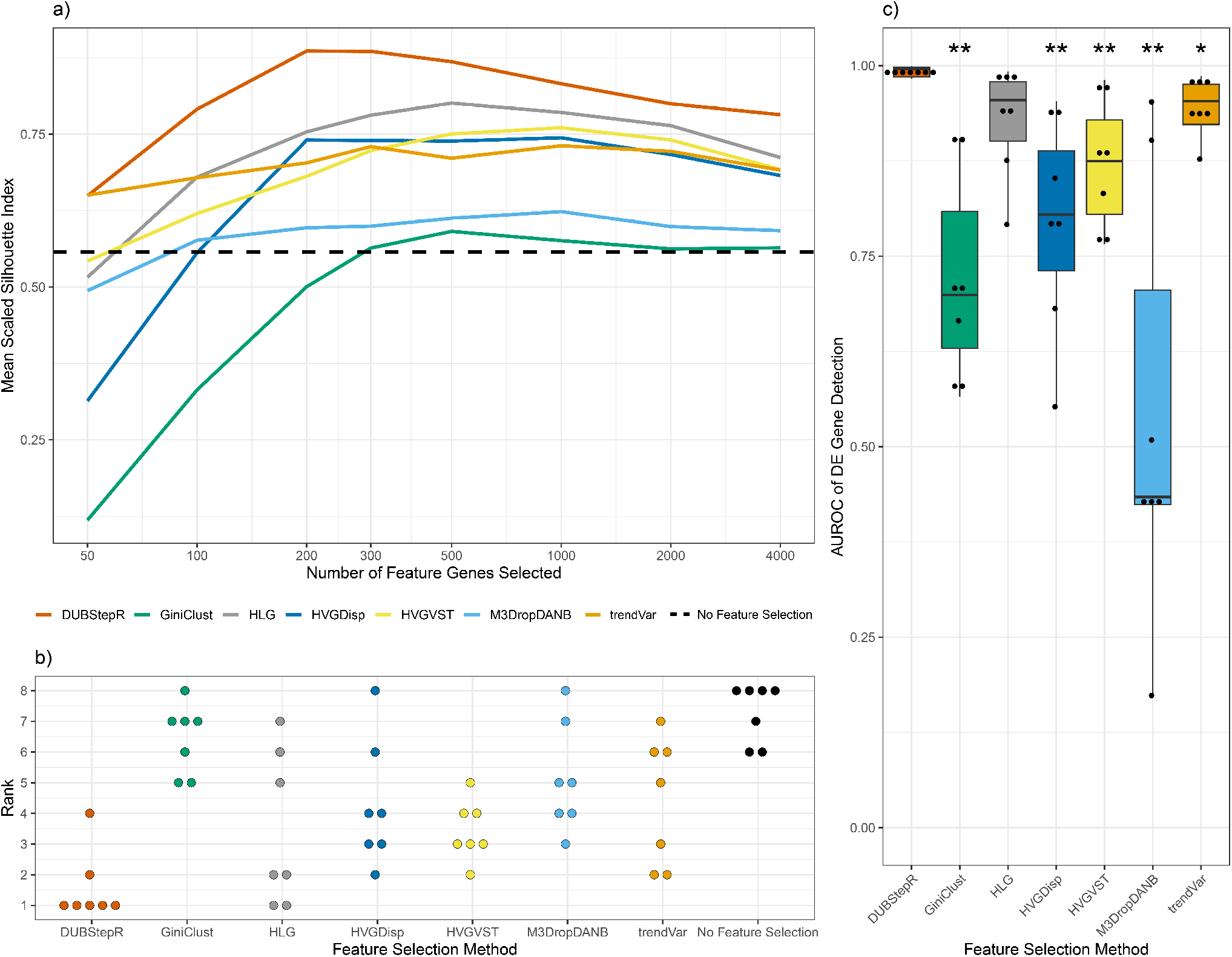
Benchmarking feature selection methods. a) Mean scaled Silhouette Index of feature sets ranging from 50 to 4, 000 features. b) Rank distribution of feature selection methods. For each dataset, the 7 methods are ranked from 1 to 7 by their best SI across all feature set sizes. c) AUROC of DE gene detection. *: p <= 0.05, **: p <= 0.01.

For optimal cell type clustering, a feature selection algorithm should ideally select only cell-type-specific genes (DE genes) as features. As an independent benchmark, we therefore quantified the ability of feature selection algorithms to discriminate between DE and non-DE genes. To minimize the effect of ambiguously classified genes, we designated the top 500 most differentially expressed genes in each dataset as DE, and the bottom 500 as non-DE (Methods), and then quantified performance using the area under the receiver operating characteristic (AUROC). Remarkably, DUBStepR achieved an AUROC in excess of 0.97 on all 7 datasets, indicating near-perfect separation of DE and non-DE genes (Figure 3c). In contrast, none of the other methods exceeded a median AUROC of 0.88. Thus, DUBStepR vastly improves our ability to select cell-type-specific marker genes (DE genes) for clustering scRNA-seq data.

With the exponential increase in the size of single-cell datasets, any new computational approach in the field must be able to scale to over a million cells. To improve DUB-StepR’s ability to efficiently process large datasets, we identified a technique to reduce a key step in stepwise regression to a single matrix multiplication, sped up calculation of the elbow point and implemented the entire workflow on sparse matrices (Methods). To benchmark scalability, we profiled execution time and memory consumption of DUBStepR, as\ well as the other aforementioned feature selection methods, on a recent mouse organogenesis dataset of over 1 million cells (21). This dataset was downsampled to produce two additional datasets of 10*k* and 100*k* cells respectively, while maintaining cell type diversity (Supp. Note 2). DUBStepR, HVGDisp, HVGVST and M3Drop were able to process the entire 1 million cell dataset, while the other three algorithms could not scale to 100*k* cells (Supp. Fig. S3). On the largest dataset, DUBStepR ranked third out of 7 tested methods in memory consumption and fourth in compute time. In terms of memory scalability, DUBStepR used 3x more memory to process the 1M cell dataset as compared to the 100*k* dataset. In contrast, HVGDisp, HVGVST and M3Drop increased their memory consumption by 12.5x. Thus, DUB-StepR is scalable to over a million cells and shows promise for even larger datasets.

### Density Index predicts the optimal feature set

As shown above, selecting too few or too many feature genes can result in sub-optimal clustering (Figure 3a). Ideally, we would want to select the feature set size that maximized cell type separation (i.e. the SI) in the feature space. However, since the feature selection algorithm by definition does not know the true cell type labels, it is not possible to calculate the SI for any given feature set size. We therefore en-deavoured to define a proxy metric that would approximately model the SI without requiring knowledge of cell type labels. To this end, we defined a measure of the inhomogeneity or “clumpiness” of the distribution of cells in feature space. If each cell clump represented a distinct cell type, then this measure would tend to correlate with the SI. The measure, which we termed Density Index (DI), equals the root mean squared distance between all cell pairs, divided by the mean distance between a cell and its *k* nearest neighbours (Methods). Intuitively, when cells are well clustered and therefore inhomogeneously distributed in feature space, the distance to nearest neighbours should be minimal relative to the distance between random pairs of cells, and thus DI should be maximal (Figure 4a,b). Empirically, we found that DI and SI were indeed positively correlated and tended to reached their maxima at approximately the same feature set size (Figure 4c). Further, for 5 out of the 7 benchmarking datasets, the feature set with the highest DI also maximized SI (Figure 4d). Since our earlier analysis only tested a discrete number of feature set sizes (Figure 3a; Supplementary Note 3), the DI-guided approach even improved on the maximum SI in 2 cases (Figure 4d). One additional advantage of the DI is that it is relatively straightforward to compute, since the numerator is proportional to the square of the Frobenius norm of the gene expression matrix (Methods). By default, DUBStepR therefore selects the feature set size that maximizes DI. The complete DUBStepR workflow is shown in Figure 5.

**Fig.4.**
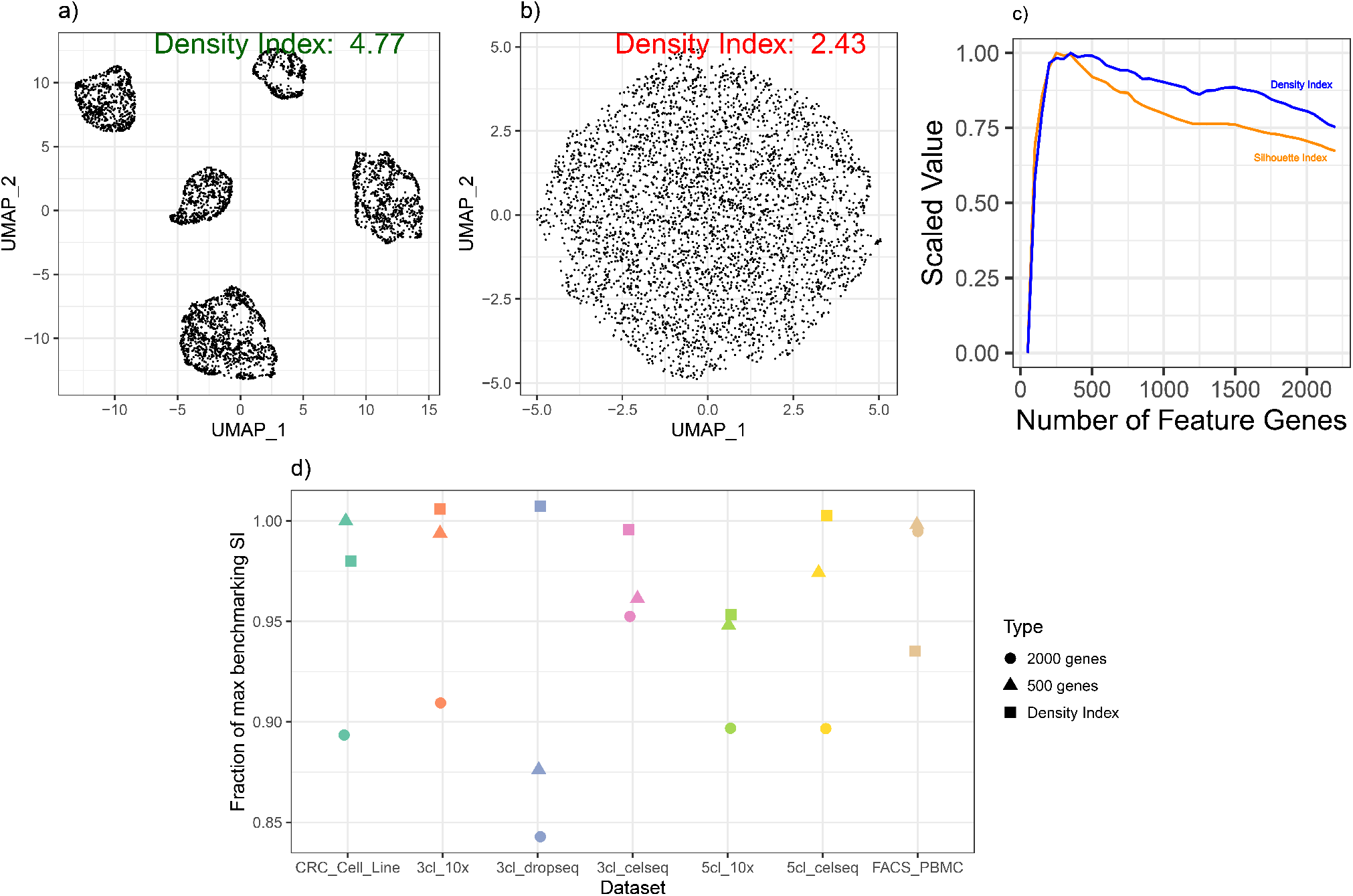
a-b) UMAP visualizations of the 5 cell lines comparing the local neighbourhood density of feature spaces in (a) good feature selection versus (b) poor feature selection. c) Comparison of optimal Density Index and Silhouette Index over feature set size range. d) Fraction of the maximum benchmarking SI achieved using feature set sizes determined by the Density Index as compared to fixing feature set sizes at 500 genes and 2000 genes. (Refer to Supp. Table S1 for details regarding the datasets.)

**Fig.5.**
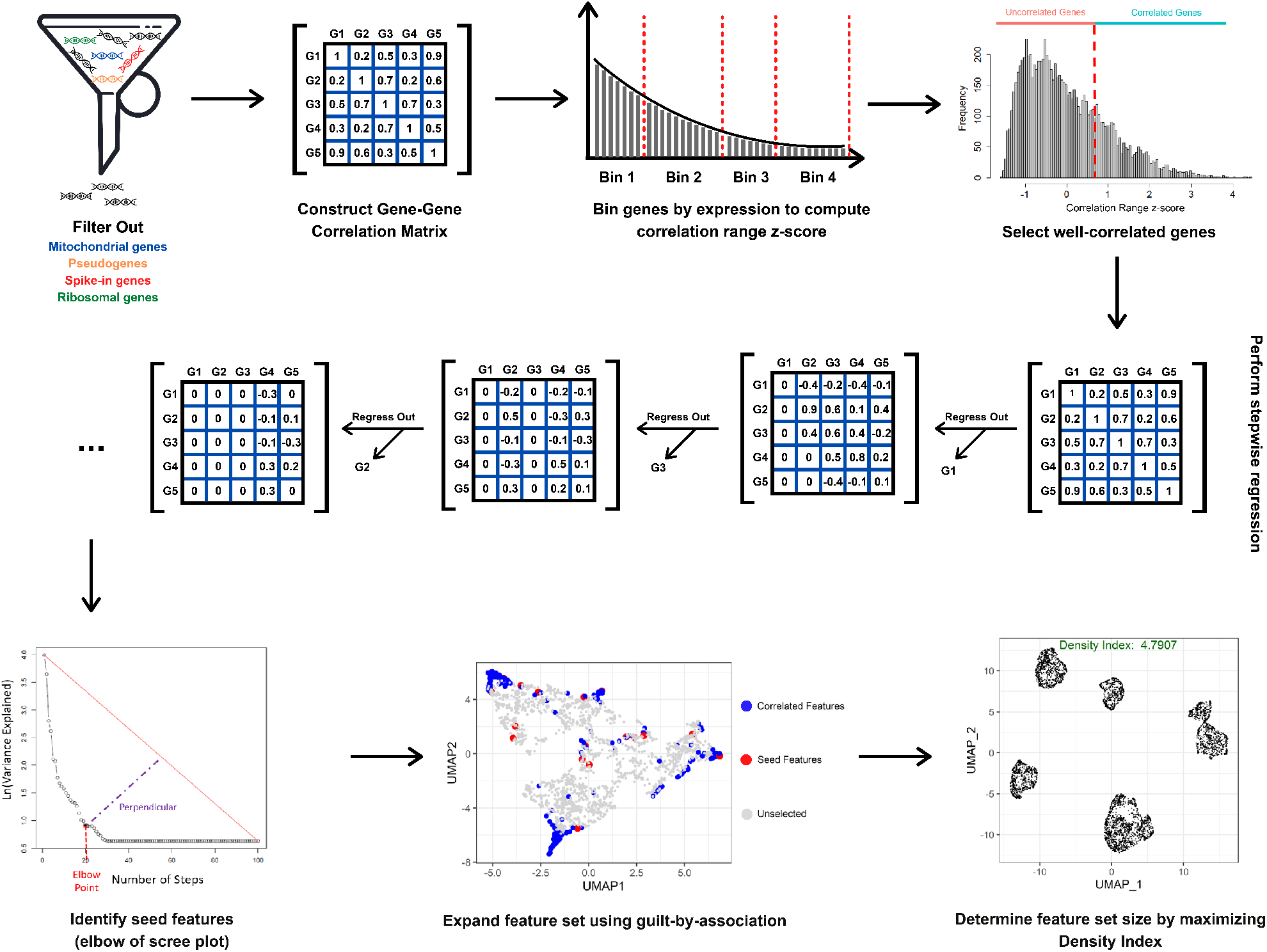
Overview of DUBStepR Workflow. After filtering out mitochondrial, ribosomal, spike-in and pseudogenes, DUBStepR constructs a GGC matrix and bins genes by expression to compute their correlation range z-scores, which are used to select well-correlated genes. DUBStepR then performs stepwise regression on the GGC matrix to identify a minimally-redundant subset of seed features, which are then expanded by adding correlated features (guilt by association). The optimal feature set size is determined using the Density Index metric.

### DUBStepR robustly clusters rare contaminating basophils in sorted PBMCs

The above benchmarking analysis was largely based on detection of relatively common cell types (> 10% of all cells). To examine the performance of feature selection methods in detecting a rare cell type, we analyzed a published scRNA-seq dataset generated from human dendritic cells (DC) and monocytes purified from PBMCs (16). In this study, the authors manually discarded 7 contaminating basophil cells based on expression of the marker genes *CLC* and *MS4A2*. To evaluate the possibility of automatically detecting this contaminating population, we applied DUBStepR to the entire dataset of 1, 085 cells. We found that DUBStepR clearly separated the 7 basophils into a distinct group (Figure 6a), which intriguingly also included 5 additional cells. We scored each cell by its average expression of the canonical basophil markers (*CCR3, CPA3, HDC, CLC, GATA2* and *MS4A2*) and found that the 5 additional cells showed high basophil marker expression, indicating that they were indeed basophils (Figure 6b,c). In contrast to DUBStepR, some of the other feature selection methods did not clearly separate this rare population of cells (Figure 6d). This result suggests that DUBStepR could also be used to detect rare cell types in scRNA-seq data.

**Fig.6.**
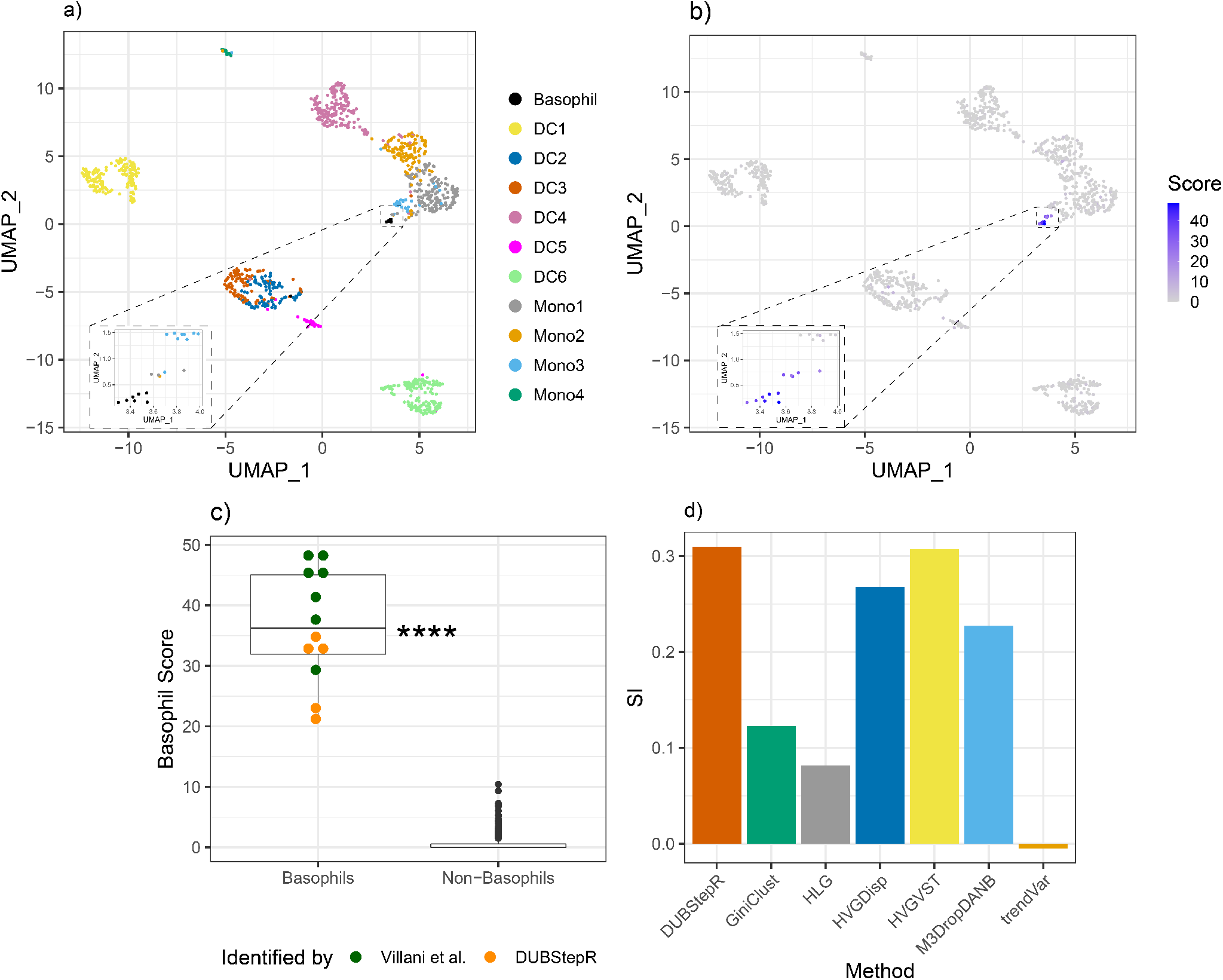
a) UMAP visualization of scRNA-seq data from Villani et al. after feature selection using DUBStepR, with cells colored by cell type labels as defined by Villani et al. Inset: zoomed-in UMAP visualization of the basophils identified by DUBStepR. b) Cumulative score of canonical basophil markers (*CCR3, CPA3, HDC, CLC, GATA2* and *MS4A2*). c) Comparison of basophil marker expression between basophils and non-basophils in the dataset. d) SI of basophils identified by DUBStepR compared to existing methods. ****: p-value <= 0.0001

### DUBStepR generalizes to scATAC-seq data

Feature selection is typically not performed on scATAC-seq data, since their mostly binary nature renders them refractory to conventional single cell feature selection based on highly variable features (23). However, since the logic of feature correlations applies even to binary values, we hypothesized that DUBStepR could also be applied to this data type. To test this hypothesis, we applied DUBStepR to scATAC-seq data from 8 FACS-purified subpopulations of human bone marrow cells (22). In contrast to the common approach of using all scATAC-seq peaks, we found that peaks selected by DUB-StepR more clearly revealed the emergence of the three ma-jor lineages from the haematopoietic stem cells: lymphoid, myeloid and megakarocyte/erythroid (Figure 7a, b). Pseudotemporal trajectories generated using Monocle 3 (21) further corroborated this result (Figure 7c, d).

**Fig.7.**
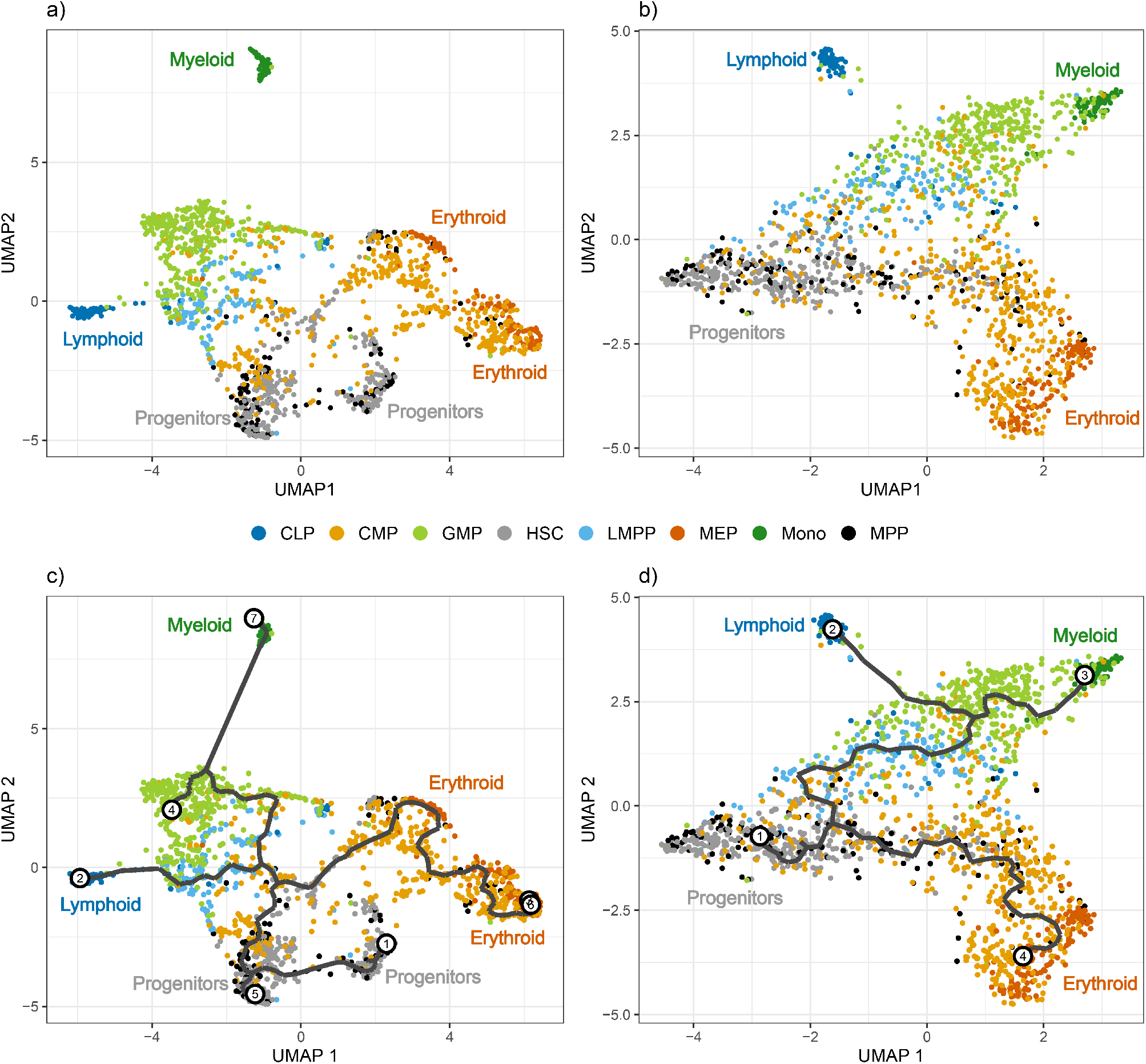
UMAP visualization of scATAC-seq data from FACS-purified hematopoietic cell populations from human bone marrow (22). a) All scATAC-seq peaks, without any feature selection. b) Peaks selected by DUBStepR. c-d) Trajectories generated by pseudotemporal ordering using Monocle 3. c) All scATAC-seq peaks, and d) Peaks selected by DUBStepR. CLP: Common Lymphoid Progenitor, CMP: Common Myeloid Progenitor, GMP: Granulocyte/Macrophage Progenitor, HSC: Haematopoietic Stem Cell, LMPP: Lymphoid-primed Multi-Potential Progenitor, MEP: Megakaryocyte/Erythroid Progenitor, Mono: Monocyte, MPP: Multi-Potent Progenitor.

## Discussion

DUBStepR is based on the intuition that cell-type-specific marker genes tend to be well correlated with each other, i.e. they typically have strong positive and negative correlations with other marker genes. After filtering genes based on a correlation range score, DUBStepR exploits structure in the gene-gene correlation matrix to prioritize genes as features for clustering. To benchmark this feature selection strategy, we used a stringently defined collection of single cell datasets for which cell type annotations could be independently ascertained (15). Note that this avoids the circularity of defining the ground truth based on the output of one of the algorithms being tested. Results from our benchmarking analyses indicate that, regardless of feature set size, DUBStepR separates cell types more clearly other methods (Figure 3a, b). This observation is further corroborated by the fact that DUBStepR predicts cell-type-specific marker genes substantially more accurately than other methods (Figure 3c). Thus, our results demonstrate that gene-gene correlations, which are ignored by conventional feature selection algorithms, provide a powerful basis for feature selection.

The plummeting cost of sequencing, coupled with rapid progress in single-cell technologies, has made scalability an essential feature of novel single cell algorithms. DUBStepR scales effectively to datasets of over a million cells without sharp increases in time or memory consumption (Supp. Fig. S3). Thus, the method is likely to scale well beyond a million cells. A major factor in the algorithm’s scalability is the fact that, once the gene-gene correlation matrix is constructed, the time and memory complexity of downstream steps is constant with respect to number of cells.

Intriguingly, DUBStepR approaches its maximum Silhouette Index value at 200-500 feature genes (Supplementary Figure S2), which is well below the default feature set size of 2,000 used in most single cell studies (10, 12). Thus, our results suggest that, if feature section is optimized, it may not be necessary to select a larger number of feature genes. Note however that the optimum feature set size can vary across datasets (Supplementary Figure S2). Selecting a fixed number of feature genes for all datasets could therefore result in sub-optimal clustering (Figure 4d).

From the perspective of cell clustering, the optimal feature set size is that which maximizes cell type separation in feature space, which can be quantified using the SI. As an indi-rect correlate of cell type separation, we have defined a measure of the inhomogeneity or “clumpiness” of cells in feature space, which we termed the Density Index (DI). To our knowledge, DI is the only metric for scoring feature gene sets based on the distribution of cells in feature space. Our results suggest that the DI correlates with the SI, and that cluster separation is improved in most cases when the feature set is chosen to maximize DI. Another important advantage of the DI is that it is computationally straightforward to calculate from the Frobenius norm of the data matrix. It is possible that the DI measure could also be applied to other stages of the clustering pipeline, including dimensionality reduction (selecting the optimal number of PCs) and evaluation of normalization strategies.

Interestingly, although DUBStepR was not specifically designed to detect rare cell types, it nevertheless showed the greatest efficacy in distinguishing the 12 contaminating basophils present in the Villani et al. dataset (Figure 6d). Notably, the original study identified only the subset of 7 basophils contained within the dendritic cell pool our analysis demonstrates that basophil contamination was also present in the sorted monocyte population (Figure 6a).

Algorithmic pipelines for single cell epigenomic data, for example scATAC-seq, typically do not incorporate a formal feature selection step (23, 24). In most cases, such pipelines merely discard genomic bins at the extremes of high and low sequence coverage. This is because the sparsity and near-binary nature of single-cell epigenomic reduces the efficacy of conventional feature selection based on mean-variance analysis. Since DUBStepR uses an orthogonal strategy based on correlations between features, it is less vulnerable to the limitations of single cell epigenomics data (Figure 7). Thus, DUBStepR opens up the possibility of incorporating a feature selection step in single cell epigenomic pipelines, including scATAC-seq, scChIP-seq and single-cell methylome sequencing.

## Code Availability

DUBStepR is freely available as an R package on GitHub at https://github.com/prabhakarlab/DUBStepR, and is well documented for easy integration into the Seurat pipeline. Code for generating all the figures in this paper is available on Zenodo at https://doi.org/10.5281/zenodo.4072260.

## Data Availability

All datasets used in this paper are publicly available, as described in Supplementary Note 2. Processed data used for generating the figures in this paper are available on Zenodo at https://doi.org/10.5281/zenodo.4072260.

## Acknowledgements

The authors would like to acknowledge Muzlifah Haniffa and Gary Reynolds for assistance with analysis of the Villani et al. dataset, Mohammad Amin Honardoost and Lavanya M. Iyer for their mentorship and guidance, and all other members of the Prabhakar lab for critical feedback and discussions. This publication is part of the Human Cell Atlas -www.humancellatlas.org/publications.

## Author Contributions

BR and SP designed the DUBStepR algorithm, with critical inputs from WS, JP, VS, IJ and FS. BR, WS and KM developed the software and performed benchmarking analyses, with assistance from JP, RX, FA, FS, NAR and MGLL. BR and SP wrote the manuscript.

## Funding

The work was supported by the Call for Data Analytics Proposal (CDAP) grant no. 1727600056.

## Methods

### Gene Filtering

By default, DUBStepR filters out genes that are not expressed in at least 5% of cells. We allow the user to adjust this parameter if they are interested in genes that are more sparsely expressed in their data set. Additionally, mitochondrial, ribosomal and pseudogenes (using the Ensembl pseudogene reference from GRCh38 (25)) are removed.

### Correlation Range

Correlation range *c*_*i*_ for gene *i* can be defined for a gene-gene correlation matrix *G* as

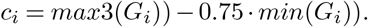

Correlation range uses the second-largest non-self correlation value (denoted here as *max*3) to calculate the range, to protect against genes with overlapping 5’ or 3’ exons (26). The minimum correlation value has been down-weighted to 0.75 to give greater importance to stronger positive correlations over negative correlations.

We first binned genes based on their mean expression level, as mean expression tends to correlate with technical noise (27). In each bin, we compute a z-score of correlation range of gene *i* as

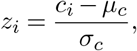

where *µ*_*c*_ is the mean correlation range of the gene and σ_*c*_ refers to variance in the correlation range scores of a gene. Genes with a z-score ≤ 0.7 are filtered out at this step.

### Step-wise Regression

We define the step-wise regression equation as

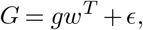

where *G* is the column-wise zero-centered gene-gene correlation matrix, *g* is the column of the matrix *G* to be regressed out, *ϵ* is the matrix of residuals and *w* is a vector of regression coefficients. The squared error (*ϵ*^*T*^ *ϵ*) is minimized when

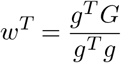

Thus,

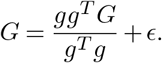

We calculate variance explained by the regression step as 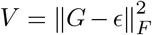, where *F* indicates the Frobenius norm. To efficiently compute *V* for all genes, we define *X* = *G*^*T*^ *G* and *x* as the row of *X* corresponding to gene *g*. Thus, *x* = *g*^*T*^ *G*. We can simplify *V* as

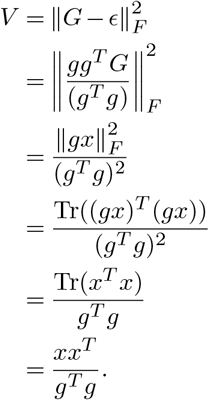

Thus, we can use a single matrix multiplication *G*^*T*^ *G* to efficiently calculate variance explained by each gene in the gene-gene correlation matrix, and then regress out the gene explaining the greatest variance. The residual from each step *k* is then used as the gene-gene correlation matrix for the next step. In other words,

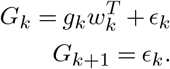

For computational efficiency, we repeat this regression step 30 times and then assume that the next 70 steps explain the amount of variance as the 30th step, giving a total of 100 steps. We observed that this shorter procedure had little or no impact on the results, since the variance explained changed only marginally beyond the 30th step.

To select the genes contributing to the major directions in *G*, we use the elbow point on a scree plot. The elbow point is the point on the scree plot that has the largest perpendicular distance to the line connecting the first and last points of the plot.

Once the genes representing the major directions of variation in gene correlations are identified using the elbow plot, they are used as seed genes to add correlated genes through guilt-by-association (28) in a single-linkage manner.

### Density Index

For a given feature set, Principal Component Analysis (PCA) (5) is performed on the gene expression matrix and the top *D* principal components (PCs) are selected, where *D* is a user-specified parameter with a default value of 20. Let *M* be the matrix of embeddings of the gene expression vectors of *N* cells in *D* principal components. The root-mean-squared distance *d*_*rms*_ between pairs of cells *i* and *j* can be calculated as

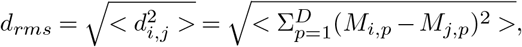

where <> denotes the average over all pairs {(*i, j*) | *i* ∈ [1, *N*], *j* ∈ [1, *N*]}. Note that, for simplicity of the final result, we include pairs in which *i* = *j*. This can be further simplified as follows:

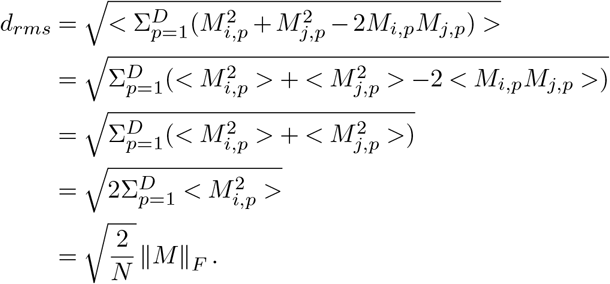

In the above derivation, the mean product term < *M*_*i,p*_*M*_*j,p*_ > is zero because *M*_*i,p*_ and *M*_*j,p*_ have zero mean across *i* and *j* respectively. Let *k*_*i*_ denote the average distance of cell *i* from its *k* nearest neighbours, and *k*_*m*_ denote the mean of *k*_*i*_ across all cells. We define the Density Index (DI) as

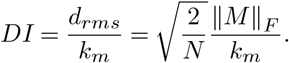

## Supplementary Note 1: Supplementary Methods

### A. Determining DE and non-DE genes

For visualization in Figure 1a-b, DE genes were determined using the FindMarkers function in Seurat using the following parameters: *logfc*.*threshold* = 1, *min*.*pct* = 0.5, *test*.*use* = ”*wilcox*”, *only*.*pos* = *T*. The top 10 DE genes of each cell line were selected for visualization.

For the rest of Figure 1, DE genes were calculated in a pair-wise fashion using the reclusterDEConsensus function as implemented in the scConsensus (29) package using the following parameters: *method* = ”*W ilcoxon*”, *meanScalingF actor* = 1, *qV alT hrs* = 0.1, *f cT hrs* = 2, *deepSplitV alues* = 1: 4, *minClusterSize* = 10. For each gene, out of all the pair-wise cell line comparisons, the largest absolute log2(fold-change) and -log10(q-value) was used. DE genes were selected as those genes whose best absolute log2(fold-change) > *log*2(1.5) and best -log10(q-value) > 1.

For Figure 1c, the 50 genes with lowest absolute log2(fold-change) values were selected as non-DE genes.

## Supplementary Note 2: Datasets

### A. Datasets for benchmarking feature selection performance

We used 5 datasets from Tian et al. (15), consisting of 3 datasets consisting of 3 cell lines each and 2 datasets consisting of 5 cell lines. In addition, we used an in-house-generated colorectal cancer (CRC) cell line dataset, first published in Li et al. (30), and FACS-sorted PBMCs from Zheng et al. (31) Figure S1 shows heatmaps of the ground truth clusters used for benchmarking feature selection performance.

### 3 cell line datasets

The H2228, H1975 and HCC827 cell lines were used to make these datasets. The 3 datasets were generated by sequencing these cell lines on CEL-Seq2, 10X Genomics and Drop-seq platforms. All datasets were processed using the `Seurat` package (v3.1.0) (10).

The 10X and Drop-seq datasets were normalized using the LogNormalize function in `Seurat`, with a scale factor of 10000 and a pseudo-count of 1, before log transformation. The CEL-Seq2 dataset was logCPM normalized i.e. its cells were normalized to a scale factor of 1000000, before log transformation with a pseudo-count of 1. Finally, the genes in these datasets were filtered so that any gene expressed in less than 5% of cells were discarded.

### 5 cell line datasets

The H2228, H1975, HCC827, H838 and A549 cell lines were used to make these datasets. The 2 datasets were generated by sequencing these cell lines on CEL-Seq2 and 10X Genomics platforms.

For the 5 cell line dataset sequenced using CEL-Seq2, we combined the data obtained from the 3 plates (p1, p2 and p3) into one single dataset. We further selected only those cells that the demultiplexing result provided predicted as single cells. Finally, the data was logCPM normalized i.e. its cells were normalized to a scale factor of 1000000, before log transformation with a pseudo-count of 1. The 5 cell line dataset sequenced using 10X was normalized using the LogNormalize function in Seurat, with a scale factor of 10000 and a pseudo-count of 1, before log transformation.

### CRC Cell Line dataset

The FPKM data from these cell lines was downloaded from GEO (GSE81861). This data has already passed the QC metrics defined by the authors. After removing replicates, we included the remaining 460 cells in this dataset.

### FACS PBMC dataset

The dataset was downloaded from the 10X Genomics website, and 2, 600 cells were sampled from CD14+ Monocytes, B cells, CD34+ Cells, Naive CD4+ T cells, Naive CD8+ T cells and NK cells each. The resulting 13, 000 cells were subjected to quality control measures. A cell would only be included in the dataset if it had greater than 300 but less than 2, 000 detected genes, and had a mitochondrial rate (proportion of reads originating from mitochondrial genes) of less than 8%. Due to sparsity of reads in this dataset, our standard filtering of genes expressed in 5% of cells rendered less than 4, 000 genes (2, 332 genes). Hence, for this dataset, we modified our filtering threshold to keep genes with expressed in at least 100 cells (approximately 1% of cells).

**Table S1.** Number of cells and genes in each dataset used for benchmarking performance of feature selection methods.

| Dataset | Num Cells | Num Genes | Abbreviation |
| --- | --- | --- | --- |
| 3 cell line CEL-Seq2 | 240 | 21,126 | 3cl_celseq |
| 3 cell line 10X | 895 | 14,479 | 3cl_10x |
| 3 cell line Drop-Seq | 210 | 13,460 | 3cl_dropseq |
| 5 cell line CEL-Seq2 | 571 | 12,616 | 5cl_celseq |
| 5 cell line 10X | 3,822 | 11,675 | 5cl_10x |
| CRC Cell Line | 460 | 32,916 | CRC_Cell_Line |
| FACS PBMC | 12,679 | 32,738 | FACS_PBMC |

### B. Dataset for benchmarking computational scalability

The Mouse Organogenesis Cell Atlas dataset was used for benchmarking scalability. The raw counts matrix for the dataset was downloaded from the Seattle Organismal Molecular Atlases (SOMA) Data portal (21). The genes were converted from Ensembl IDs to gene symbols for compatibility with DUB-StepR using `biomaRt` (32, 33). The counts were first used to form a Seurat Object, which was downsampled to roughly 10, 000 and 100, 000 cells using the `subset` function in `Seurat` (v3.1.0). Cluster labels provided by Cao et al. were used to downsample and ensure that all cell types were equally represented in the datasets, although rarer cell types were fewer in number. We opted to downsample the same dataset to preserve biological heterogeneity, and to allow benchmarking comparability. Following this, the dataset was converted to a SingleCellExperiment object. The dataset was filtered to retain genes expressed in at least 1 cell and log-transformed with a pseudocount of 1, using the *logNormCounts* function within the `scuttle` R package. The log-transformed counts matrix was used as the input for the various algorithms tested.

### C. Villani et al. dataset

The TPM expression matrix was downloaded from GSE94820. We followed the quality control (QC) metrics as stated in Villani et al., i.e. we only included cells having over 3000 detected genes. Further, genes were filtered so that only genes expressed in at least 0.5% of the cells were kept in the dataset.

After QC, the data was log-transformed with a pseudocount of 1, and the log-TPM data was used as input for DUBStepR. DUBStepR was run on this data using default settings, except for the change in gene filtering. Due to the presence of rare clusters in this dataset, we modified the gene filtering criterion to filter out genes expressed in less than 1% of cells. After feature selection, the features were scaled and centered, and principal component analysis was run using the Seurat package. 15 PCs were chosen for Silhouette Index computation. The Seurat package was also used to generate the UMAP and feature plot visualizations in Figure 6.

### D. Single-cell ATAC sequencing dataset

We obtained the scATAC-seq dataset of human hematopoietic progenitors from Buenrostro et al (22). The data was provided after processing, and provided as a peaks-vs-cells matrix, wherein the peaks had been selected from the bulk hematopoietic ATAC-seq data in the paper.

All peaks that were not accessible in at least 100 cells were removed from the dataset. This resulted in a data matrix of 2, 034 cells with 32, 090 peaks. Finally, every cell was normalized to a scale factor of 1000 using read-count normalization in Seurat, so as to account for differences in overall accessibility of the cells. This normalized data was fed into DUBStepR for feature selection.

After feature selection, PCA was performed on both the normalized data without feature selection and the dataset with DUBStepR-selected peaks. 50 PCs were used to capture the variance in the dataset in both cases, and the output of PCA was used for UMAP visualization.

For pseudotemporal ordering using Monocle 3, the `Seurat` object with the UMAP coordinates was converted to a Monocle *cell_data_set* object. Cells were clustered in Monocle 3 in the UMAP space using the *cluster_cells* function, with the following parameters: *resolution* = 1*e* − 3 and *num*_*iter* = 10.

## Supplementary Note 3: Benchmarking

### A. Silhouette Index computation

To compare feature selection methods in terms of cell type separation, we modified a commonly used metric known as Silhouette Index (17). The Silhouette Index is a measure of how close a cell is to other cells of the same cluster. Since we know the ground-truth cell type assignments for our cells in the datasets, we can compute the Silhouette Index for each cell. Typically, the Silhouette Index of a feature space is the average of the Silhouette Index values for each cell. However, we noted that this manner of computing Silhouette Index was disadvantageous to rare cell types, as clusters with fewer cells would have a lower contribution to the average.

Thus, we modified the calculation of Silhouette Index to give equal weight to each cell type in the dataset. We first calculate the average Silhouette Index value for cells in each cluster, and take the mean of those values to compute the Mean Silhouette Index (Mean SI). This way, regardless of its abundance, each cell type is given equal weight. Cell-cell distances are calculated in the principal component space after selecting the top 20 PCs.

**Fig.S1.**
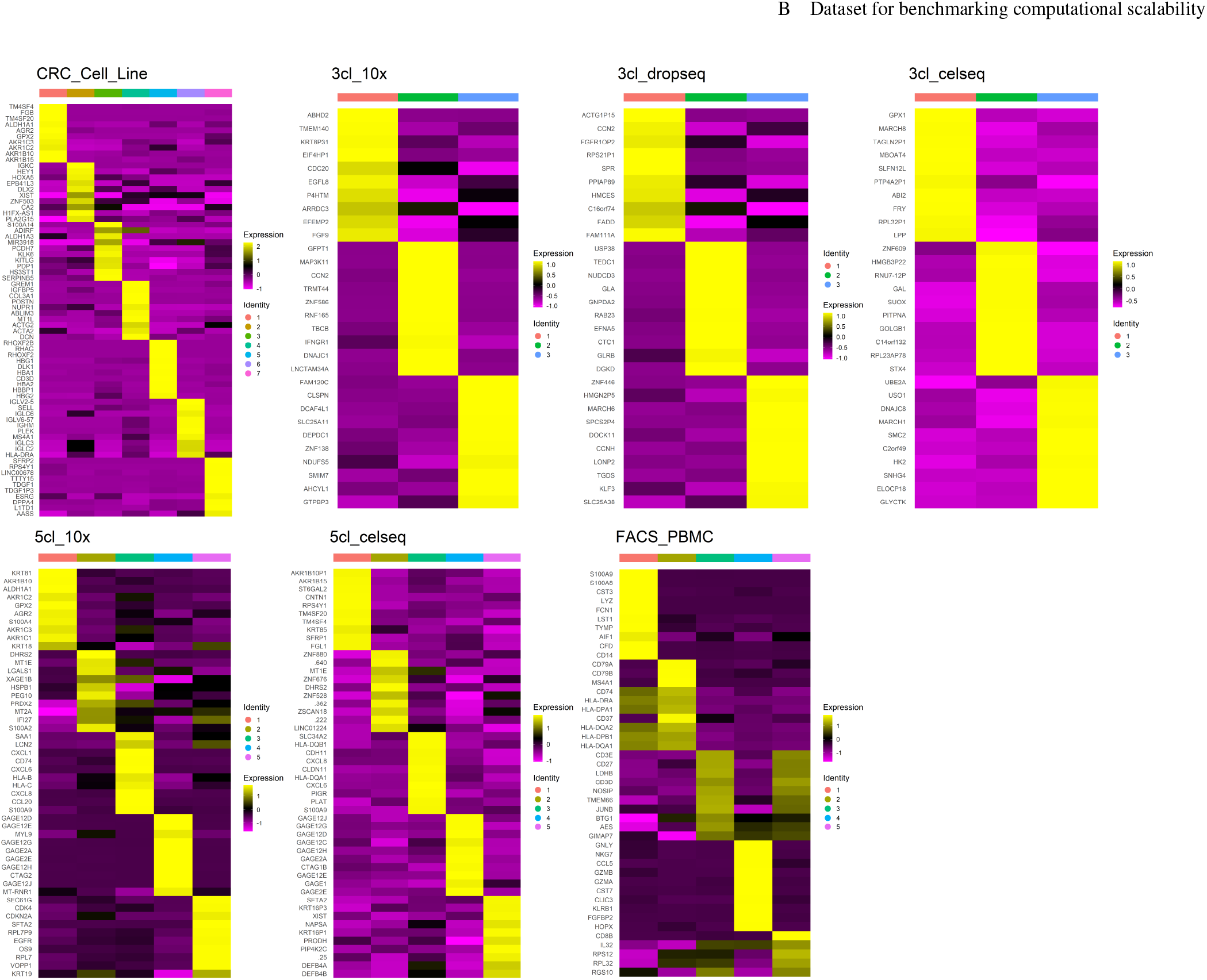
Heatmaps showing ground truth of data sets used for benchmarking feature selection algorithms.

### B. AUC of marker gene detection

For the computation of AUC, genes in these datasets were filtered so that any gene expressed in less than 5% of cells were discarded. For each dataset, the top 500 genes were selected based as marker genes and the bottom 500 genes as non-marker genes based on their best q-value of pair-wise cluster-specific differential expression. For each feature selection method, genes were ordered based on the score provided by the method. The feature selection algorithm and corresponding ordering metric are explained in Table S3. The AUC was computed as the area under the Receiver Operating Characteristic (ROC) curve i.e. the curve between the true positive rate (TPR) and false positive rate (FPR) in the selection of marker genes.

**Table S2.**
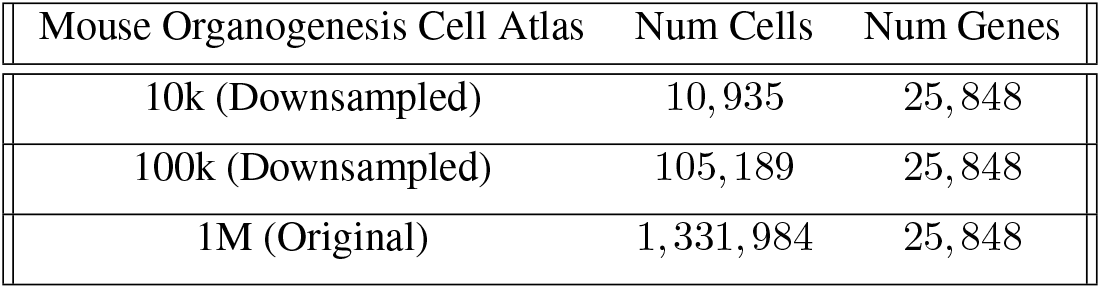
Number of cells and genes in each dataset used for benchmarking computational scalability of feature selection methods.

### C. Execution time and memory consumption calculation

To benchmark scalability of feature selection algorithms, we used the *Rprof()* function from R `utils` v3.6.2 to profile memory consumption and execution time. To obtain both time and memory, we set the parameter *memory* = ”*both*” to additionally report the change in total memory. We converted time from milliseconds into minutes by dividing the output time by 60000. In case of memory usage, we converted the output memory from Mb to GB by dividing the output memory by 8000. Profiling was done on our lab’s Ubuntu server with 64 cores (clock speed 2.2GHz) and 1.5 TB of RAM.

**Table S3.** Metrics used to order genes output by the benchmarked feature selection algorithms.

| Method | Ordered by |
| --- | --- |
| DUBStepR | Guilt-by-association order, then correlation range |
| HVGDisp | dispersion |
| HVGVST | variance stabilized transformation (vst) |
| M3Drop/DANB | p-value |
| trendVar | biological component |
| GiniClust | p-value |
| HLG | PC contribution scores |

## Supplementary Note 4: Supplementary Figures

**Fig.S2.**
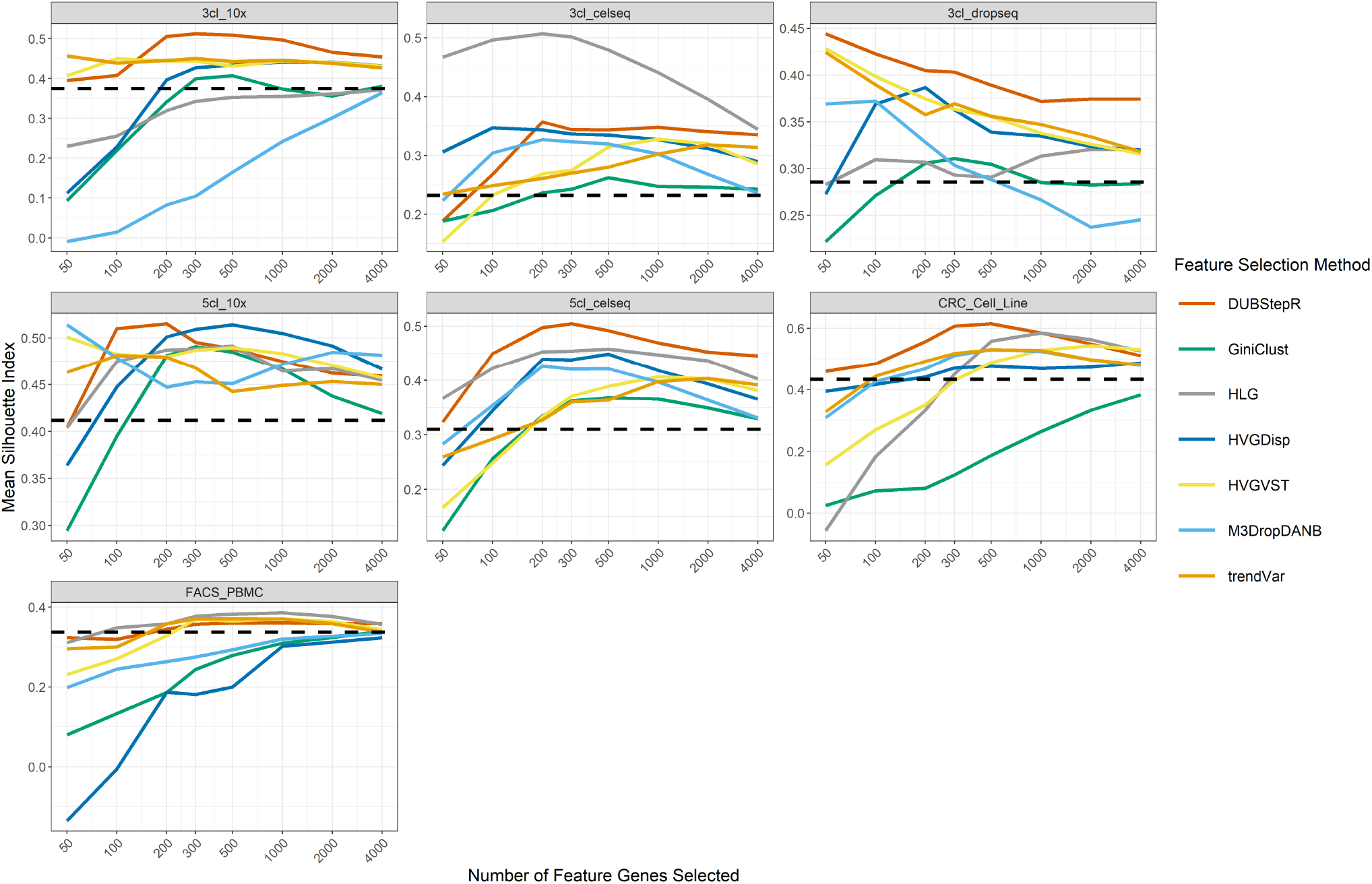
Line plots of SI over different numbers of feature genes selected. Black dashed line indicates SI without feature selection.

**Fig.S3.**
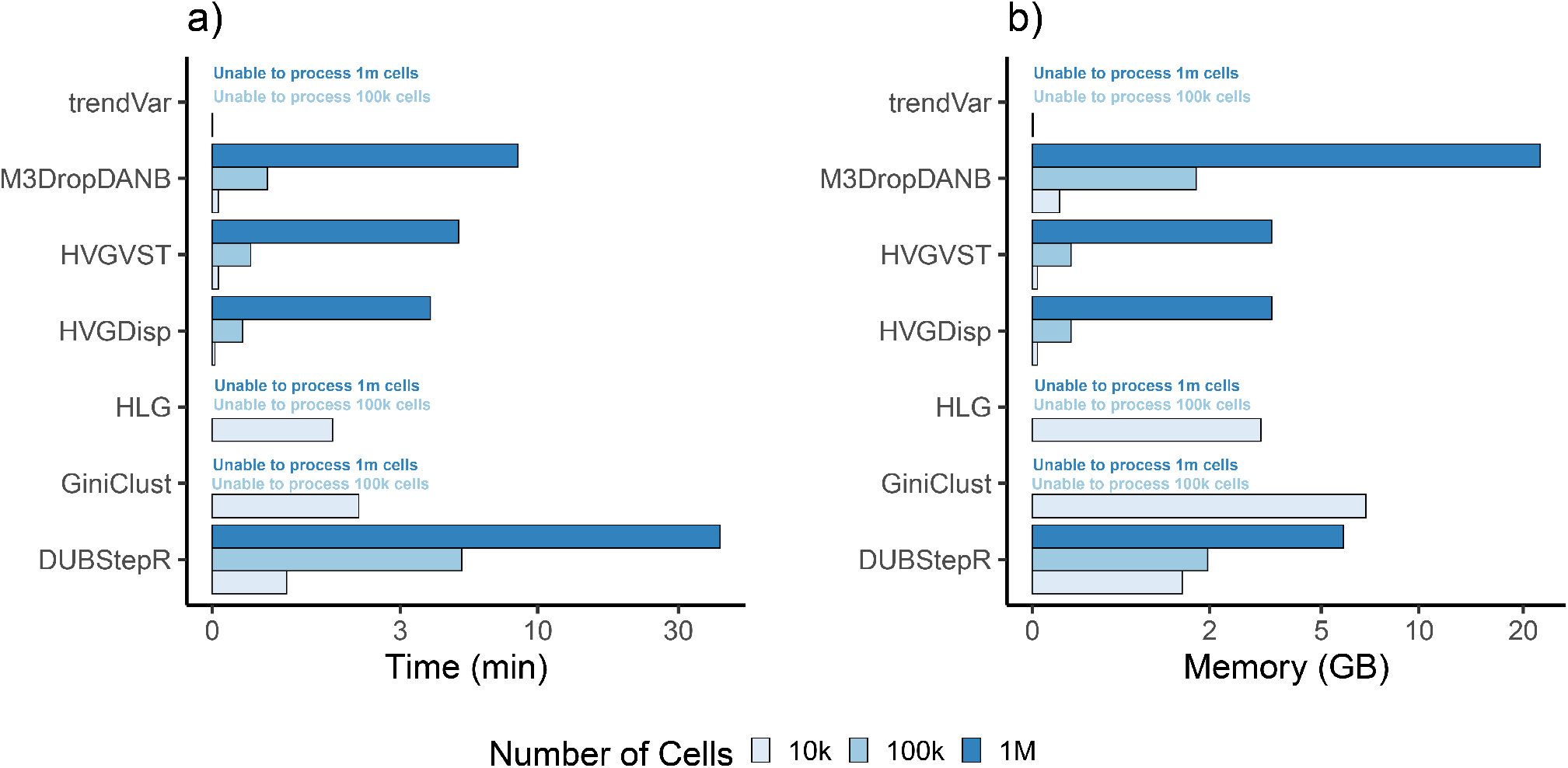
Benchmarking of computational efficiency of DUBStepR against existing feature selection methods on datasets of 10*k*, 100*k* and 1 million cells. a) Execution time (in minutes) taken of each method. b) Total memory consumed (in GB) by each method. The X-axes of both plots were log-transformed for ease of visualization.

**Fig.S4.**
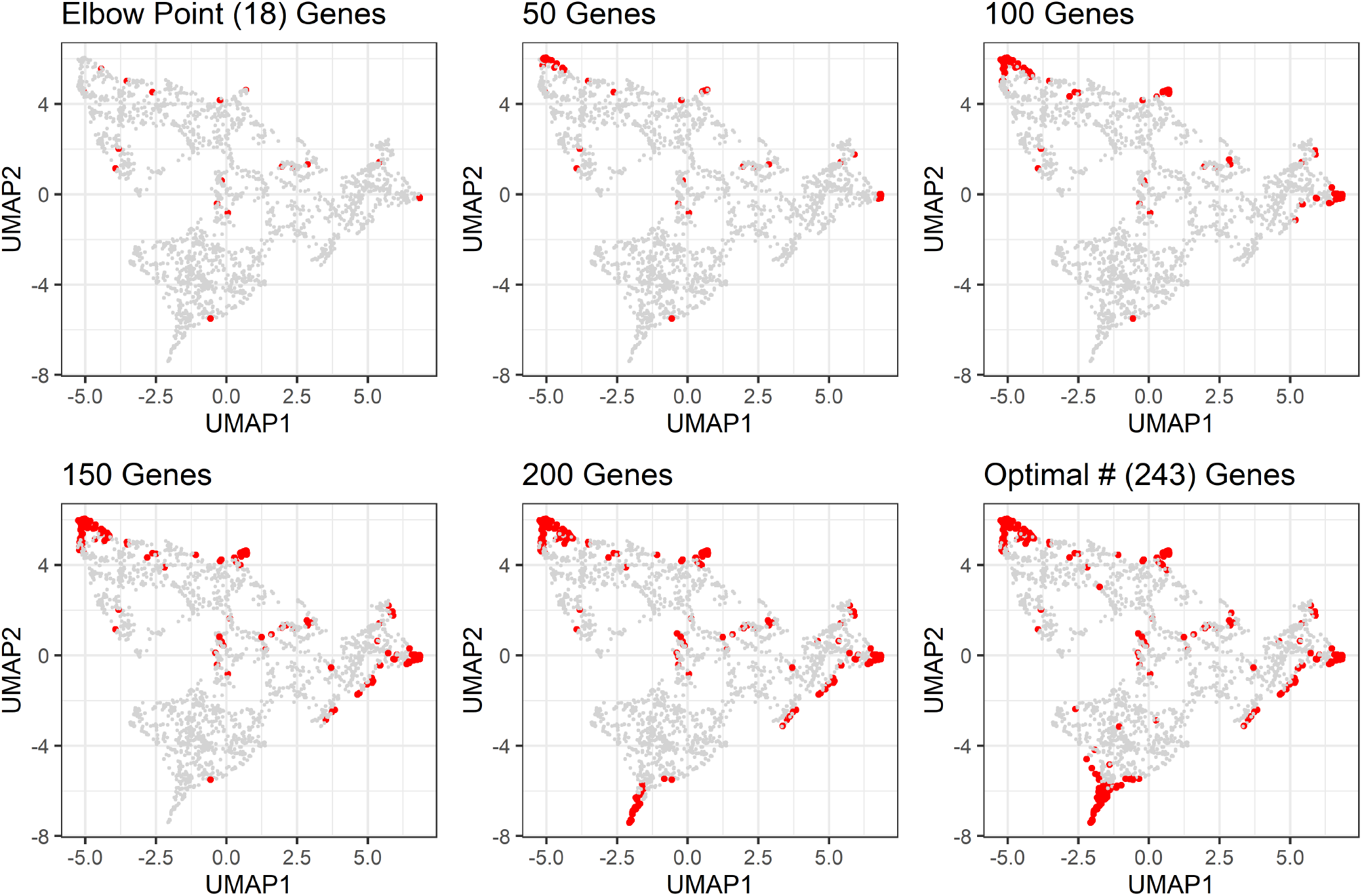
UMAP visualization of the gene-gene correlation matrix. Red: Genes selected as features. Grey: Genes not selected as features.

## Notes

https://github.com/prabhakarlab/DUBStepR

https://doi.org/10.5281/zenodo.4072260

### Competing Interest Statement

The authors have declared no competing interest.

### Summary of Updates

Updated with editor's suggestions.

